# A unifying framework for structured population dynamics in modern coexistence theory

**DOI:** 10.64898/2026.11.13.766574

**Authors:** Joe Brennan

**Affiliations:** Department of Evolution and Ecology, University of California, Davis, USA 95616; Center for Population Biology, University of California, Davis, USA 95616

## Abstract

Variation within populations is ubiquitous in nature, yet many models and frameworks in community ecology assume that all individuals within a population interact with the community identically. Modern coexistence theory is no exception; however, several subfields of ecology have independently been integrating coexistence theory, each asking the same underlying question: how do structured population dynamics shape coexistence outcomes? Here, I propose a unifying framework for integrating population variation into modern coexistence theory that is applicable across all fields of ecology. The application of this framework is presented in three case studies: niche overlap and fitness differences between influenza substrains, mutual invasibility between competing perennial plants, and fluctuation-dependent invasion outcomes in apparently competing prey with a foraging-predation risk tradeoff. Together, these examples illustrate the framework’s broad applicability, power to resolve the interaction between population and community level processes, and its potential to reveal novel ecological phenomena.

## Introduction

Within-population variation is ubiquitous (Des Roches et al., 2018). The resulting population structure can impact community dynamics through multiple axes of variation, including age and stage (Miller and Rudolf, 2011), genotype and phenotype (Crutsinger et al., 2006; Post et al., 2008), behavior (Fryxell and Lundberg, 2012; Gil et al., 2018), space (Binckley and Resetarits, 2005; García-Callejas et al., 2019), and disease status (Hudson and Greenman, 1998; Hatcher et al., 2006). The role of intraspecific variation on community dynamics is so pervasive that it is often proposed as a mechanism for resolving the persistent mismatch between predicted and observed coexistence outcomes (Nakazawa, 2015; Uriarte and Menge, 2018). Despite mounting evidence, coexistence theory has developed largely under the assumption that such variation is negligible and can therefore be ignored (Bolnick et al., 2011; Violle et al., 2012).

One framework of community ecology rapidly evolving to accommodate structured population dynamics is modern coexistence theory (MCT). MCT is a quantitative framework of community ecology that uses the invasion growth rate, defined as the average per-capita growth rate of a species when rare and resident species are coexisting, to predict coexistence outcomes and understand the relative effects of different ecological factors (Chesson, 1994; Barabás et al., 2018; Ellner et al., 2019). Traditionally, MCT has been applied to models that do not explicitly model structured population dynamics. Despite this, MCT has greatly deepened our understanding of the mechanisms that promote coexistence, including fluctuation-dependent mechanisms such as relative nonlinearity (differential nonlinear responses to fluctuating ecological factors due to Jensen’s inequality) and the storage effect (differential responses to the environment & covariance between nonadditively interacting environmental conditions and competition) (Chesson, 1994). Eager to extend these ideas to communities with structured population dynamics, several studies have applied MCT to various areas of population biology where structured populations play a key role, including evolutionary ecology & game theory (Yamamichi et al., 2022; Park, 2026), demography & population ecology (Song et al., 2021; Werner et al., 2025), spatial ecology (Chesson, 2000; Shoemaker and Melbourne, 2016; Ellner et al., 2022b), and epidemiology & disease ecology (Sieben et al., 2022; Park et al., 2024).

Although these extensions have effectively leveraged MCT within their respective subdisciplines, each has developed largely in isolation, with its own formalization and notion of what “structure” means. As a result, there is no general framework for incorporating structured population dynamics into MCT, even though these studies all ask the same fundamental question: how does population structure influence coexistence? The challenge of structured population dynamics in MCT across all of these studies is the same. Calculating invasion growth rates is straightforward for unstructured populations whose dynamics are described by a single equation. However, when a population is structured, its dynamics are often described by multiple equations e.g., separate equations for distinct phenotypes within a population to account for differences in their demographic rates or selective pressures. This complicates not only the derivation of the invasion growth rate but also the expression of the per-capita growth rate in a form conducive for invasion growth rate decompositions. A unifying framework would provide a common methodology, allow intraspecific variation to be incorporated into coexistence theory regardless of how it arises, and offer a shared language for synthesizing results across ecological systems.

Here, I propose a unifying framework for incorporating structured population dynamics into MCT, allowing synthesis across subdisciplines of ecology. To do this, I first demonstrate how equations describing structured populations can be transformed to simply evaluate invasion growth rates (Patel and Schreiber, 2018; Zou et al., 2023; Kim, 2024), a necessary ingredient of MCT. Then, I demonstrate three applications of how these invasion growth rates integrating structured dynamics can be used to derive key metrics of MCT. First, I analytically derive niche overlap and fitness differences between two strains of influenza with cross immunity studied in Yang et al. (2020) and Park et al. (2024). I then demonstrate how the mutual invasibility criteria can be used to determine coexistence outcomes when niche overlap and fitness differences are difficult to derive, using a model of competing perennial plants with explicitly modeled seed and adult stages, studied in Uricchio et al. (2019) and Song et al. (2021). Lastly, I demonstrate how to evaluate the influence of fluctuations (e.g., relative nonlinearity and storage effect *sensu* Ellner et al. (2019)) on species’ invasion growth rates using a model of apparent competition between two prey species exhibiting a foraging-predation risk tradeoff. These applications demonstrate that this framework is generally applicable to many ecological disciplines concerned with structured population dynamics and that MCT is a powerful tool for understanding mechanisms of the coexistence of structured populations.

## Methods

Let there exist two interacting sets of variables: population densities *x*(*t*) = (*x*_1_(*t*), …, *x*_*m*_(*t*)) and another set of dynamic variables *y*(*t*) = (*y*_1_(*t*), …, *y*_*n*_(*t*)), known as auxiliary variables. Auxiliary variables can represent a variety of biologically relevant processes, including environmental conditions, trait dynamics, or, of particular interest here, population structure. Following Patel and Schreiber (2018), a broad class of ecological models can be written as

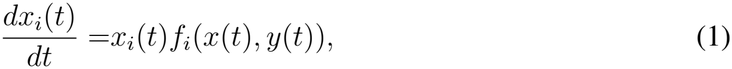

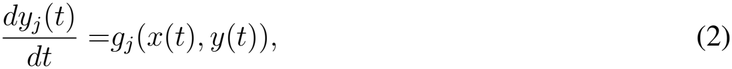

for continuous time systems or

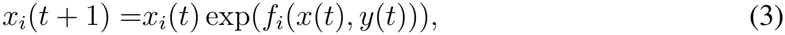

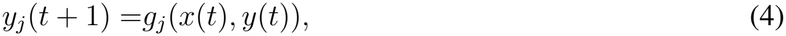

for discrete time systems where *f*_*i*_ is the per-capita growth rate of species *i* and *g*_*j*_ is a function determining how auxiliary variable *j* changes over time.

In these models, coexistence of a subset of species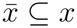, together with auxiliary variables 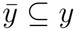, is defined by the existence of an invariant set that is bounded away from extinction and attracts all trajectories with initial conditions satisfying *x*_*i*_(0) *>* 0 for all 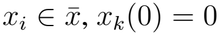, *x*_*k*_(0) = 0 for all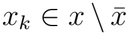, and any 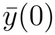 in its domain. That is, there exists a global attractor bounded away from extinction for all species in 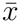, while species in 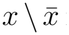 remain extinct, known as coexistence in the sense of permanence (Schreiber, 2000; Patel and Schreiber, 2018; Clark et al., 2026).

Note that auxiliary variables 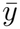 are not strictly required to be positive on this invariant set, unlike population densities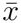.

Let *S* denote the coexistence state supporting species 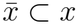 and auxiliary variables 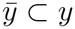.

The invasion growth rate of species *i* at coexistence state *S* is then given by

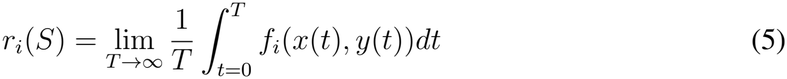

for continuous time systems or

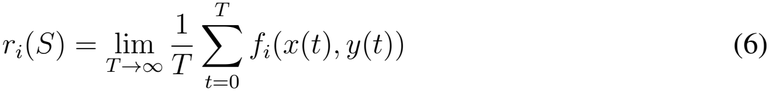

for discrete-time systems when the limit exists. If *r*_*i*_(*S*) *>* 0, species *i* will grow away from extinction in the long term. If *r*_*i*_(*S*) *<* 0, species *i* will eventually decay to extinction upon introduction at low density. Note that generally, if species *i* is supported by coexistence state *S, r*_*i*_(*S*) = 0.

To calculate the invasion growth rate of an invading species *i* using Equation 5 or 6, the function *f*_*i*_(*x*(*t*), *y*(*t*)) for *x*_*i*_(*t*) at low density (i.e., as *x*_*i*_(*t*) *→* 0) must exist. However, sometimes models of structured population dynamics are not in the form presented in Equations 1-2 or 3-4. As an illustrative example, consider an exponentially growing population consisting of non-reproductive juveniles (*N*_1_ = *N*_1_(*t*)) and reproductive adults (*N*_2_ = *N*_2_(*t*)). Immature individuals mature at rate *τ* and die at rate *d*_1_, while mature individuals produce immature offspring at rate *b* and die at rate *d*_2_, producing

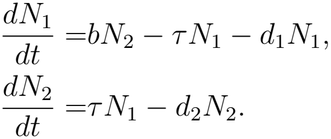

If our goal is to compute the invasion growth rate of species *x* using Equation 5, the current formulation is not suitable because it describes the dynamics of stage classes rather than species density, providing no clear per-capita growth rate function *f* . I therefore perform a change of variables that separates total population density from stage structure. Let *x* = *N*_1_ + *N*_2_ denote total population density and *y* = *N*_2_*/*(*N*_1_ + *N*_2_) denote the proportion of mature individuals. Then, I can rewrite *N*_1_ as *x*(1 *− y*) and *N*_2_ as *xy*. To define the model under this change of variables, I differentiate the equations for *x* and *y*. This gives 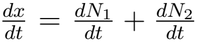 and, using the quotient rule of differentiation, 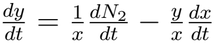 where 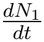 and 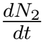 are written as a function of *x* and *y*. Under this transformation, I have

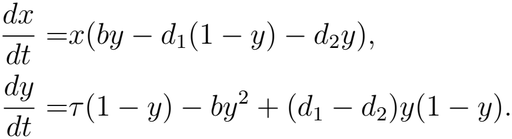

Given all parameters are positive, *y* converges to the stable equilibrium *y*^*\**^ from any initial condition 0 *≤ y*(0) *≤* 1, so the invasion growth rate is well defined and can be calculated using Equation 5 (Appendix A). This is analogous to computing the growth rate of a matrix population model under the assumption that the population has reached its stable stage distribution (Caswell, 2001). Indeed, the invasion growth rate with equilibrated stage structure equals the dominant eigenvalue of the original matrix model (proven generally in Patel and Schreiber (2018) with this specific case shown in Appendix B). More generally, when a structured population of interest is not of the form presented in Equations 1-2 and 3-4, one can perform a change of variables to separate total population density from population structure to allow ease in calculating the invasion growth rate. Specifically, the total population density can be expressed in the form of Equation 1 or 3, while the variables describing the stage structure of the population can be expressed in the form of Equation 2 or 4. This transformation allows invasion growth rates to be defined at the level of the population of interest while retaining the underlying dynamics of stage structure.

By performing this transformation in the context of community dynamics, standard analyses from modern coexistence theory can be applied. In particular, to determine whether a species with structured dynamics can invade a resident community, I allow the resident community and all associated auxiliary variables to reach their coexistence state. Simultaneously, the auxiliary variables describing the invading species are allowed to converge to the state expected when the species is rare, as visualized in Figure 1. The invasion growth rate is then calculated using Equations 5 or 6.

**Figure 1:**
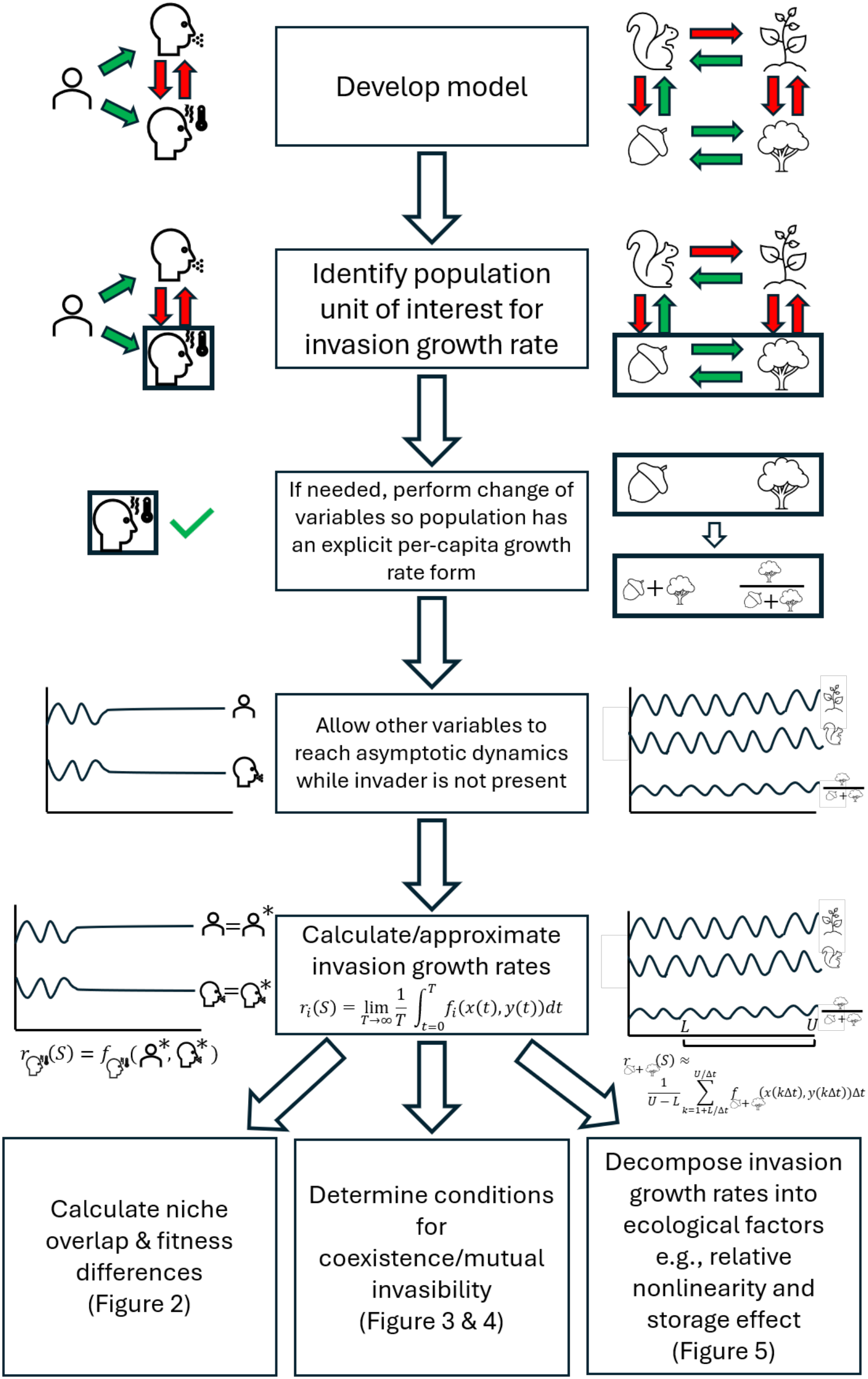
Workflow for integrating stage structure into MCT. Middle column displays general workflow while left and right columns visualize each step for a given scenario.

## Applications

To demonstrate the process of integrating structured population dynamics into modern co-existence theory (MCT), I provide three applications, each highlighting a different, core analysis used in MCT. For each case study, I work through the workflow presented in Figure 1. First, I extend Park et al. (2024) by analytically solving for niche overlap and fitness differences in a two-strain, influenza model with cross immunity. Next, I extend Song et al. (2021) by solving for prevalence of coexistence, priority effects, and competitive exclusion between two competing perennial plant species previously studied by Uricchio et al. (2019), in addition to identifying the role of perenniality in coexistence outcomes. Lastly, I study fluctuation-dependent invasion success in a model of apparently competing species with a shared predator exhibiting a foraging-predation risk tradeoff, demonstrating fluctuating population structure can in part drive a storage effect.

### Deriving niche overlap and fitness differences in a two strain influenza model with cross immunity

Park et al. (2024) proposed a framework for using MCT, specifically niche overlap and fitness differences, to understand coexistence of competing strains of disease. One of the models Park et al. (2024) studied was an extended model from Yang et al. (2020), studying the dynamics of two strains of influenza, H1N1 and H3N2, with cross-immunity. Park et al. (2024) used the model to numerically estimate niche overlap and fitness differences to predict coexistence outcomes. Here, I use my proposed framework to analytically solve for niche overlap and fitness differences between H1N1 and H3N2.

For each strain *i ∈ {*1, 2*}*, the model tracks susceptible and infected host classes, denoted *S*_*i*_ and *I*_*i*_, respectively. Susceptible hosts are replenished through births at rate *µ*, infected hosts recover at rate *γ*_*i*_, and immunity wanes at rate *δ*_*i*_. Transmission of strain *i* occurs at rate *β*_*i*_, while *σ*_*ij*_ denotes the strength of cross-immunity imposed by strain *j* on strain *i*. I assume all parameters are positive. The resulting model is

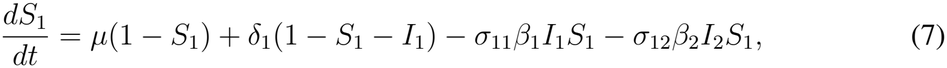

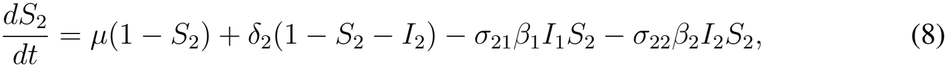

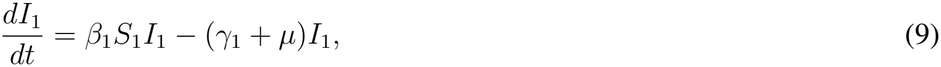

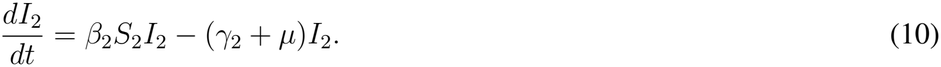

Here, we are interested in predicting coexistence outcomes of H1N1 and H3N2, with four possible outcomes: H1N1 competitively excludes H3N2, H3N2 competitively excludes H1N1, H1N1 and H3N2 coexist, or a priority effect occurs wherein the competitive exclusion outcome is contingent upon initial frequency of infected individuals in each class. Since we are interested in coexistence outcomes of only the infected classes, we denote them as our populations of interest. Thus, to use techniques from MCT, we must calculate the invasion growth rate of *I*_1_ and *I*_2_ using Equation 5.

Since both infected classes’ per-capita growth rates can be expressed as

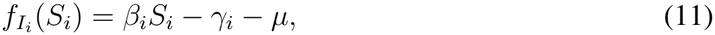

we can calculate the invasion growth rate as *I*_*i*_ approaches 0 while other populations are at some coexistence state. Thus, in this scenario, no change of variables is required to calculate the invasion growth rates using Equation 5. For this model, differential equations describing dynamics of the infected classes correspond to the form of Equation 1 while susceptible classes correspond to Equation 2.

Niche overlap and fitness differences are specific quantities that assess coexistence via mutual invasibility, where each absent strain is able to successfully invade the other strain’s persistence equilibrium. To derive niche overlap and fitness differences for the model, we must first determine the invasion growth rates of each infected class *I*_*i*_ while other populations are at their coexistence state. Precisely, we need to find 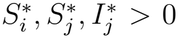 such that all Equations 7-10 evaluated at the equilibrium 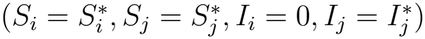 simplify to zeroes.

I first derive the invasion condition for *I*_2_ when rare and others are at some coexistence state. Let 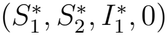 denote the equilibrium in which strain 1 persists and strain 2 is absent. The corresponding equilibrium is unique and exists when *β*_1_ *> γ*_1_ + *µ*. Hereafter, we assume this inequality holds. The equilibrium is then given by

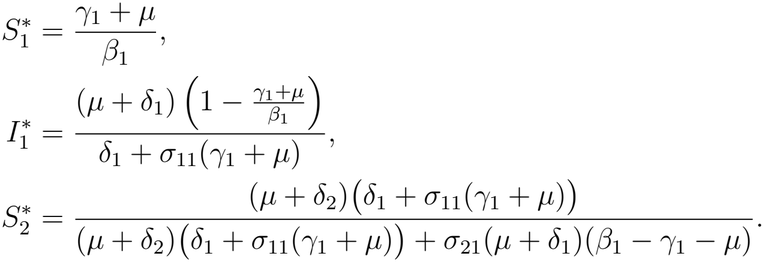

For strain 2 to successfully invade, its invasion growth rate given in Equation 11 must be positive. Substituting the equilibrium expression for 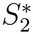 condition gives into Equation 11 and rearranging this

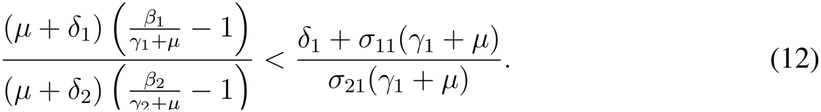

Through the same procedure, the invasion condition for strain 1 into the equilibrium 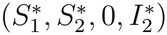 is

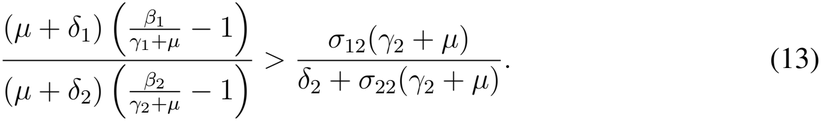

Combining equations 12 and 13 yields the mutual invasibility criterion

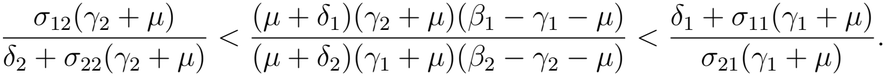

Following classic derivations of niche overlap and fitness differences, I normalize the equation by the geometric mean of the upper and lower bounds,

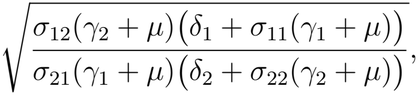

which gives

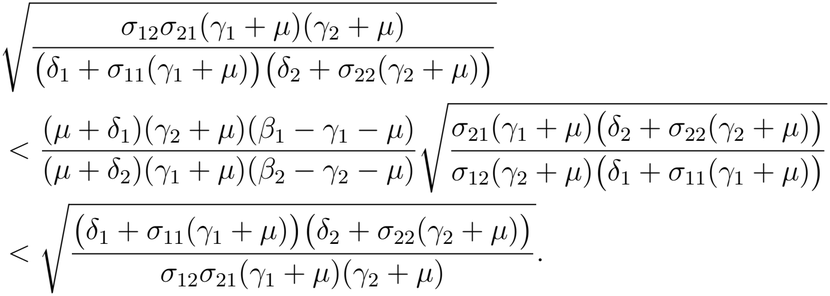

Let niche overlap be defined as

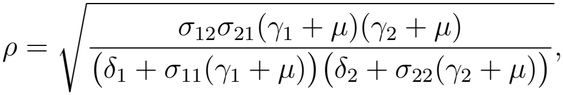

and let fitness differences be defined as

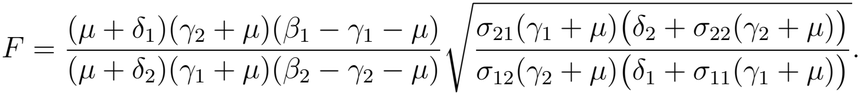

Then, we recover the classic coexistence relationship relating niche overlap to fitness differences. If 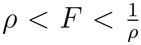, the two strains successfully coexist. If 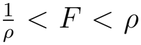, then priority effects arise. Otherwise, competitive exclusion occurs.

Now, using the parameterization presented in Appendix C studied in Yang et al. (2020) and Park et al. (2024) (though, assuming no seasonal fluctuations), I study how coexistence outcomes change as *σ*_12_ increases, modulating the competitive effect of H3N2 on H1N1 through cross immunity. I find that, under the empirically-realized parameterization, the two strains are capable of coexistence (Figure 2A & B). However, as the impact of cross immunity from H3N2 on H1N1 increases, I find that niche overlap increases and H3N2 gains larger relative fitness differences to H1N1, ultimately driving competitive exclusion of H1N1 (Figure 2A & C).

**Figure 2:**
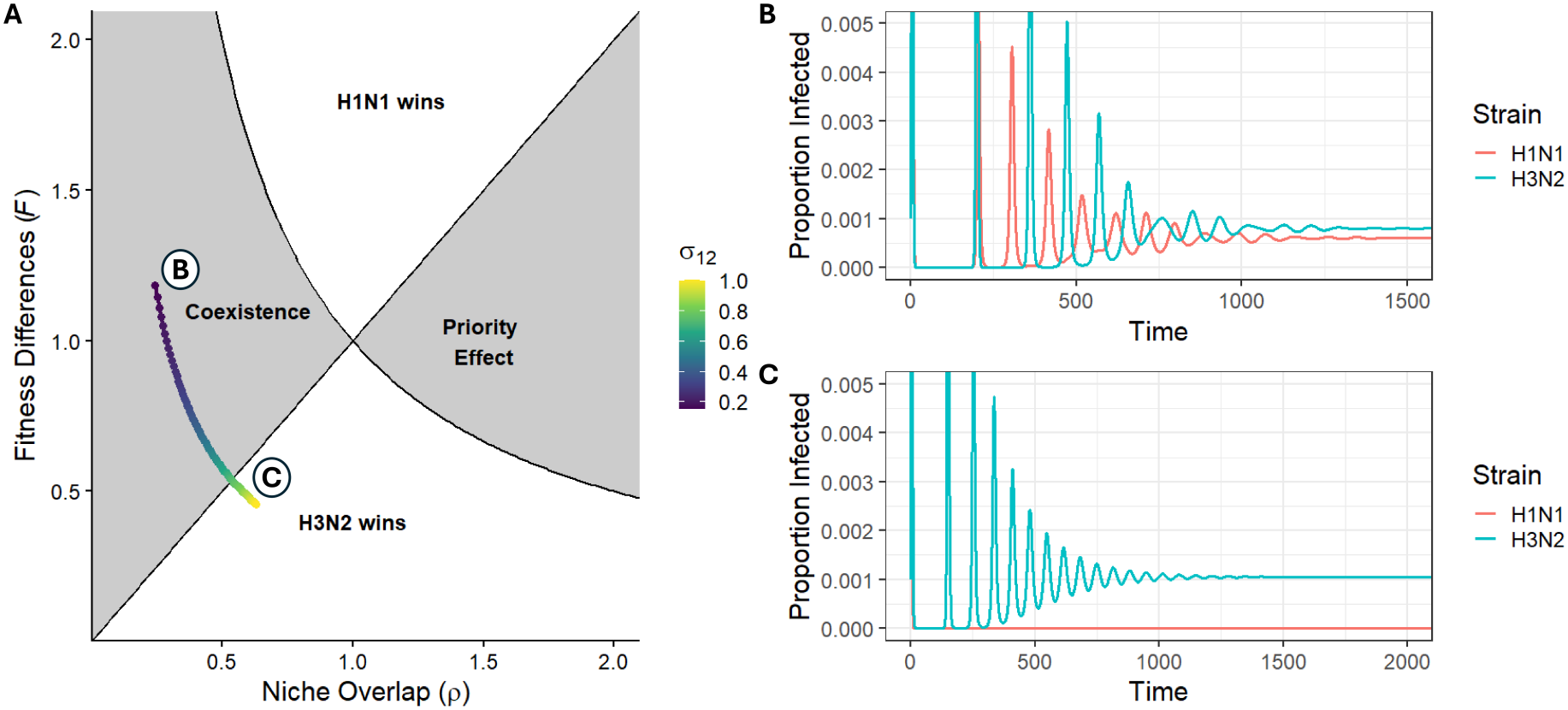
(A) Coexistence outcomes between H1N1 and H3N2 as cross immunity from H3N2 on H1N1 varies. Each region denotes a different coexistence outcome. Color of dot denotes the value of cross immunity and its placement denotes the respective niche overlap and fitness differences value. (B) Time series of infection dynamics when *σ*_12_ = 0.17, where coexistence occurs. (C) Time series of infection dynamics when *σ*_12_ = 0.75, where competitive exclusion occurs.

### Coexistence outcomes of perennial plants

Uricchio et al. (2019) studied a competitive system of native perennial plants and invasive annual plants to investigate predicted coexistence outcomes. Using this system, Song et al. (2021) explored pairwise coexistence outcomes between annual and perennial plants by deriving niche and fitness differences under some mathematical simplifying assumptions and assessed how life history influences coexistence outcomes. However, Song et al. (2021) did not evaluate coexistence outcomes between the two native perennial plants. Here, I extend this analysis by investigating predicted coexistence outcomes between the two perennial plants, *Elymus glaucus* (EG) and *Stipa pulchra* (SP), and evaluate how life history mediates these outcomes using the presented framework.

I consider a system of two competing perennial plant species. Each population is structured into two life stages: seedlings (*s*_*i,t*_ = *s*_*i*_(*t*)) and mature individuals (*m*_*i,t*_ = *m*_*i*_(*t*)). Seed production occurs exclusively from mature individuals, with a maximum per-capita fecundity given by *λ*_*i*_. This fecundity is reduced by intra- and interspecific competition, modeled through coefficients *α*_*ij*_, which capture the effect of germinated individuals of species *j* on species *i*. Seeds that are not immediately germinated are assumed to survive and carry over to the next year. Mature individuals survive from one year to the next with probability *ω*_*i*_. A fraction *g*_*i*_ of seeds germinates each year, and a maximum proportion *v* of germinated seedlings successfully survive the summer, independent of species identity. This post-germination survival is also reduced by competition from newly emerged seedlings and adults, given by *β*_*S*_ and *β*_*M*_, respectively. Under these assumptions, Song et al. (2021)’s model is then

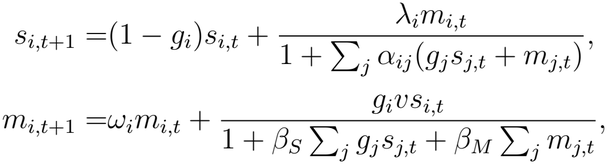

with the parameterizations provided in Appendix D.

Here, we are interested in using invasion growth rates to determine coexistence outcomes between the two species. However, to calculate the invasion growth rate through Equation 6, we need the model in the form of Equation 3 & 4. Thus, I perform a change of variables such that, instead of tracking the abundance of seedlings and mature individuals, I track the total number of individuals (*n*_*i,t*_) of the population and the proportion of those individuals that are mature (*p*_*i,t*_).

Let *n*_*i,t*_ = *s*_*i,t*_ + *m*_*i,t*_ and 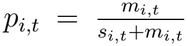. Note that, under this change of coordinates, *s*_*i,t*_ = (1 *− p*_*i,t*_)*n*_*i,t*_ and *m*_*i,t*_ = *p*_*i,t*_*n*_*i,t*_. Thus, the transformed model is

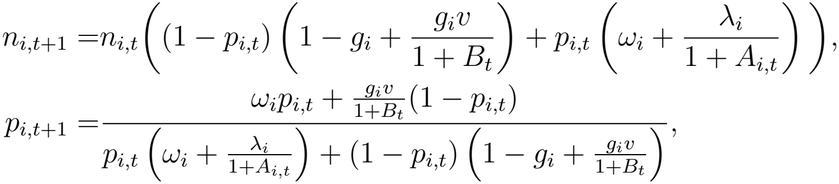

where *A*_*i,t*_ = Σ _*j*_ *α*_*ij*_(*g*_*j*_(1*−p*_*j,t*_)*n*_*j,t*_ +*p*_*j,t*_*n*_*j,t*_) and *B*_*t*_ = *β*_*S*_ *Σ* _*j*_ *g*_*j*_(1*−p*_*j,t*_)*n*_*j,t*_ +*β*_*M*_ *Σ* _*j*_ *p*_*j,t*_*n*_*j,t*_. Under this change of coordinates, the discrete-time equation describing total abundance of species *i* corresponds to Equation 3 and the equation corresponding to proportion of species *i* that is mature corresponds to Equation 4. Thus, we now can now calculate invasion growth rates using Equation 6.

The classic technique for evaluating coexistence in MCT is the mutual invasibility criterion, positing that a pair of species can coexist if each can invade the other’s single-species equilibrium. For mutual invasibility to identify coexistence outcomes correctly, the dynamics must be bounded (i.e., the system is dissipative), and each single-speces persistence state (i.e., every invariant set on the extinction boundary) must be repelling (Roth et al., 2017).

As in many models of competition, the dynamics of this model are bounded: at high density, negative density dependence in intraspecific competition causes the population to decline in the next time step such that trajectories remain bounded above while staying nonnegative. To identify single-species persistence states, I ran 100 simulations of 2000 time steps for each parameterization, each starting from different initial conditions, and examined the final 500 time steps to characterize the long-term dynamics. For all parameterizations studied, each species in monoculture persisted at a stable equilibrium, and trajectories converged to the same equilibrium regardless of initial conditions. I therefore conclude, based on numerical simulations, that each species has a single globally attracting positive equilibrium for each parameterization. Because the model has bounded dynamics and a globally attracting positive equilibrium for the parameters of interest, invasion growth rates evaluated at the single-species equilibrium determine coexistence outcomes, as shown in Figure 3.

**Figure 3:**
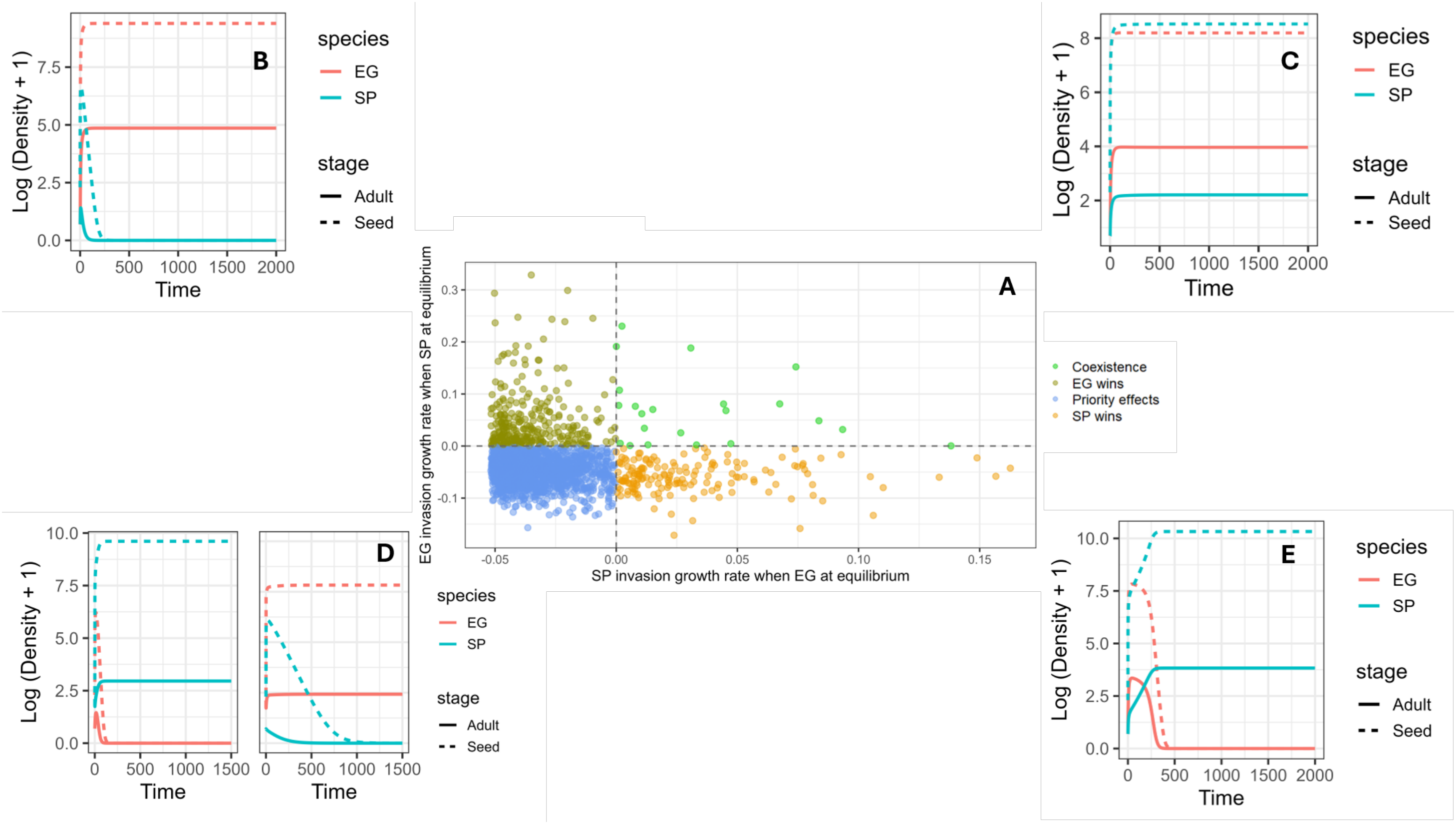
(A) Coexistence outcomes across 2000 different parameter draws. Color of points denote coexistence outcomes of a parameter draw. Dotted lines denote when axes equal zero. (B) Time series of community dynamics from parameter draw where EG outcompetes SP. (C) Time series of community dynamics from parameter draw where SP and EG coexist. (D) Time series of community dynamics from parameter draw where priority effect arise. (E) Time series of community dynamics from parameter draw where SP outcompetes EG.

There are four possible coexistence outcomes: coexistence, SP competitively excludes EG, EG competitively excludes SP, or priority effects. For coexistence to occur, we must demonstrate mutual invasibility; that is, both the invasion growth rate of EG and SP must be positive when the other species is at equilibrium (Figure 3C). Competitive exclusion of species *i* occurs if species *j* can invade the equilibrium of species *i* but species *i* cannot invade the equilibrium of species *j* (Figure 3B,E). If both species have negative invasion growth rates at the other species equilibrium, then priority effects occur (Figure 3D). To assess the most likely coexistence out-come between EG and SP using invasion growth rates, I evaluate invasion growth rates across 2000 parameterization samples from the joint posterior distribution (Song et al., 2021).

Extending analyses done in Song et al. (2021), I assess coexistence outcomes when both populations have stage structure, only one has stage structure, or when neither have stage structure. To do this, we follow a similar procedure as Song et al. (2021), removing over-summer survival of mature individuals (*ω*_*i*_ = 0), over-summer maturation of seedlings (*v* = 0) and the associated competition terms for over-summer maturation (*β*_*S*_ = *β*_*M*_ = 0). Additionally, we assume that the competition of an annual species on over-summer maturation on a perennial plant is given by a different parameter *β*_*A*_, empirically estimated in Song et al. (2021). With this, we then only track the number of seeds year to year, as adults without perenniality never survive to the following generation. The equations used to study these scenarios are shown in Appendix E.

To estimate invasion growth rates numerically, I initialized the simulation with one species and ran its dynamics for 2000 time steps. Because most parameterizations reached their stable equilibrium within about 500 time steps, I discarded the first 500 as transient, retained the last 1500, and calculated the per-capita growth rate of the absent species at each retained step. I then approximated its invasion growth rate as

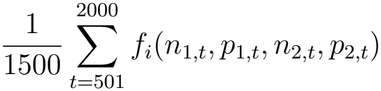

where the function *f* is defined by Equation 3.

I find most parameterizations where both species have perenniality predict priority effects (*∼* 76%) with the next most likely outcome being competitive exclusion of SP (*∼* 15%) (Figure 3). However, when I remove perenniality from one of the species, the species without perenniality becomes increasingly likely to competitively exclude the other (*∼* 97% and *∼* 94% for EG and SP without perenniality, respectively) (Figure 4). When perenniality is removed from both species’ dynamics, I find that priority effects again become the most likely coexistence outcome (*∼* 79%) (Figure 4).

**Figure 4:**
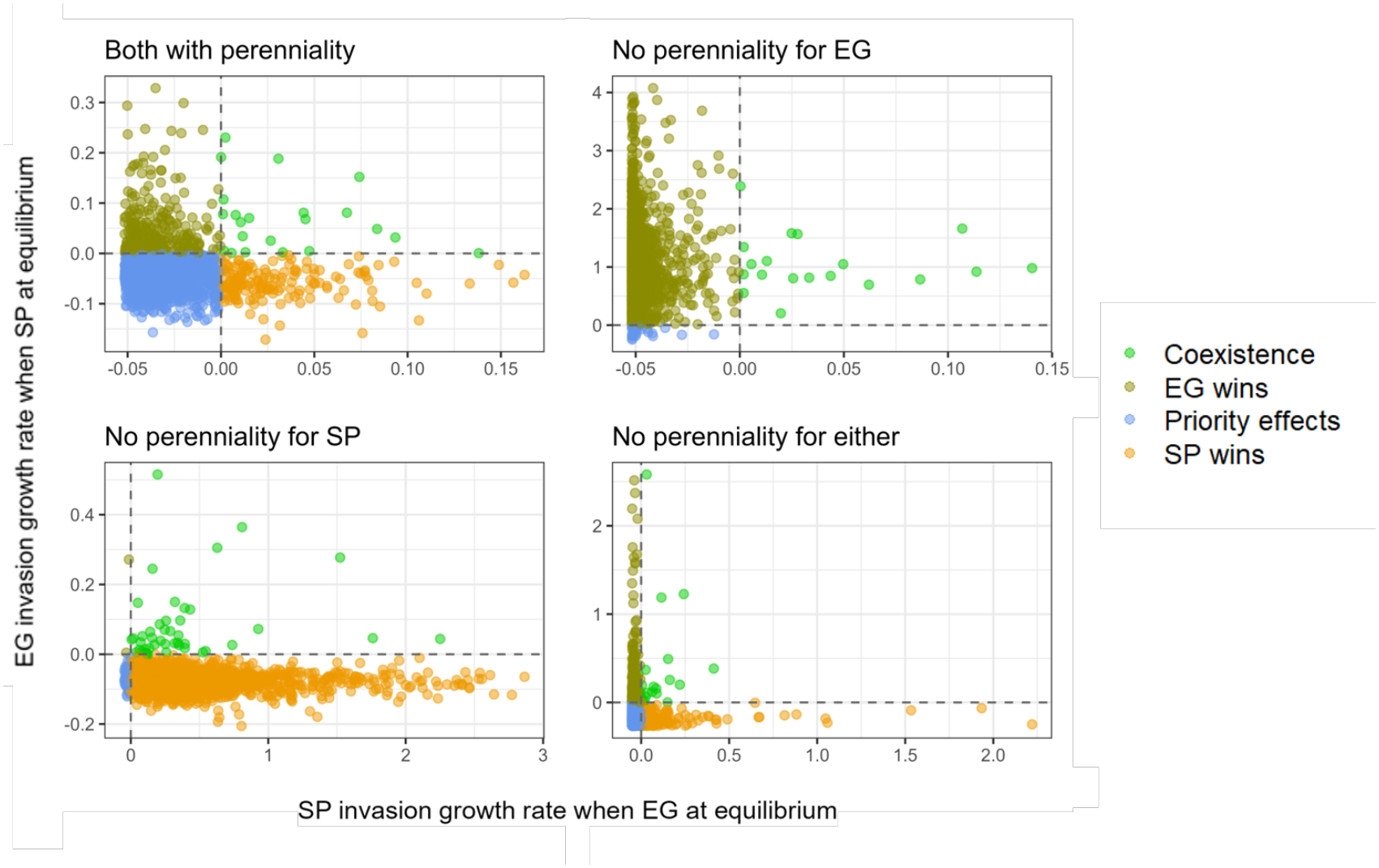
(A) Coexistence outcomes from parameter draws when both species retain stage structure. (B) Coexistence outcomes from parameter draws when EG has no stage structure. (C) Coexistence outcomes from parameter draws when SP has no stage structure. (D) Coexistence outcomes from parameter draws when neither species has stage structure.

Fluctuation dependent invasion outcome under foraging-predation risk trade-off Many species face a tradeoff wherein foraging is necessary for growth but leaves species increasingly susceptible to predation, known as the foraging-predation risk tradeoff (Sih, 1980; Abrams, 1984; Verdolin, 2006). Here, I use the proposed framework to investigate how fluctuations in predator abundance and the proportion of actively foraging individuals in a resident prey population impact the invasion success of an invading prey population under the foraging-predation risk tradeoff.

I assume there exist two species experiencing apparent competition under a shared predator (Holt, 1977). Let each species *i ∈ {*1, 2*}* have two behavioral classes: idle and foraging.

Individuals in the idle class do not contribute to population growth but are less susceptible to predation with an attack rate of *α*_*IP,i*_. Conversely, foraging individuals in species *i* contribute to population growth by converting resources into biomass/new individuals at some maximum rate *r*_*i*_; however, they face higher predation pressure with an attack rate *α*_*F P,i*_ *> α*_*IP,i*_. I assume that species *i* spends an average amount of time *τ*_*IF,i*_ in the foraging class and an average time of *τ*_*F I,i*_ in the idle class. The predator converts some proportion *c* of consumed biomass into predator biomass and has a species-independent handling time of *h*. Under these assumptions, the model is given by

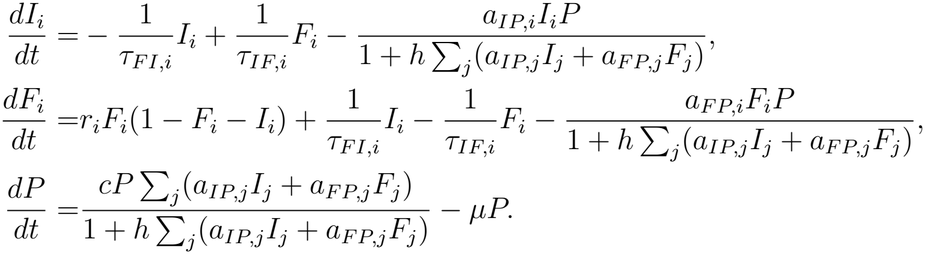

Here, we are interested in understanding if prey species 2 can invade when prey species 1 and the predator are coexisting. Under the current model formulation, we are unable to evaluate the invasion growth rate of the invading prey species using Equation 5. Thus, I define a change of variables. Let *x*_*i*_ = *I*_*i*_ + *F*_*i*_ and 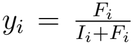. Then, I can convert our variables of interest by the equations *I*_*i*_ = *x*_*i*_(1 *− y*_*i*_) and *F*_*i*_ = *x*_*i*_*y*_*i*_. To define the new differential equations, I differentiate the new variables to arrive at 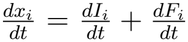 and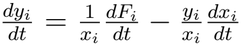, where 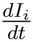 and 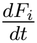 is a function of variables in our new coordinate system. Under this new coordinate system, we have

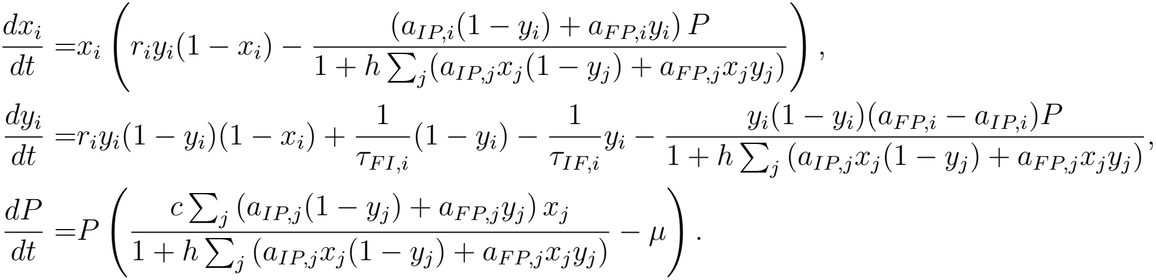

Under this formulation, the differential equation describing total prey species density corresponds to Equation 1 while the differential equation corresponding to the proportion of the population foraging corresponds to Equation 2. Thus, the invading prey population’s invasion growth rate can be calculated using Equation 5. Now with the calculated invasion growth rate, I can decompose the invasion growth rate of prey species 2 to investigate the role of fluctuations on invasion success. Specifically, I investigate the role fluctuations in predator abundance *P* and fluctuations in the proportion of the resident prey population actively foraging *y*_1_. I also calculated the impact of the interactions between the fluctuating factors, notably the impact of their covariance called the storage effect *sensu* Ellner et al. (2019). *To do this, I use the decomposition method proposed in Ellner et al*. *(2019)* described briefly in Box 1 and described in mathematical detail in Appendix F.

To numerically estimate the invasion growth rate of the invading prey population, I numeri-*cally integrated the model using the deSolve* package in R evaluated at time intervals Δ*t* = 0.1 for total run time of 20, 000 (Soetaert et al., 2010). Using this time series, I approximate the invasion growth rate of invading prey population *x*_2_ by

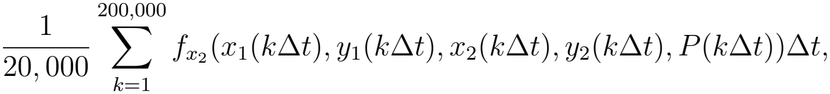

where the function *f* is defined by Equation 1.

I find that, for the parameterization studied in Appendix G, the invading species is unable to successfully invade the community (Figure 5C). Through the decomposition in Figure 5A, I find that fluctuations have significant impacts on the invasion success of the invading prey population. First, I find that fluctuations in predator density weakened predation pressure for the invading species, leading to a positive impact on the invasion growth rate of the invading prey population. Conversely, fluctuations in the resident prey population’s stage structure has a negative impact on the invading species’ invasion growth rate by weakening the effect of handling time of the predator on the invading species. Interestingly, the most influential factor on the invasion growth rate of the invading prey population was the storage effect, wherein the covariance between predator density and the proportion of foraging individuals in the resident prey population drive a strong, negative impact. This effect is strong enough such that, in the absence of the storage effect, the decomposition would predict successful invasion of the invading species.

**Figure 5:**
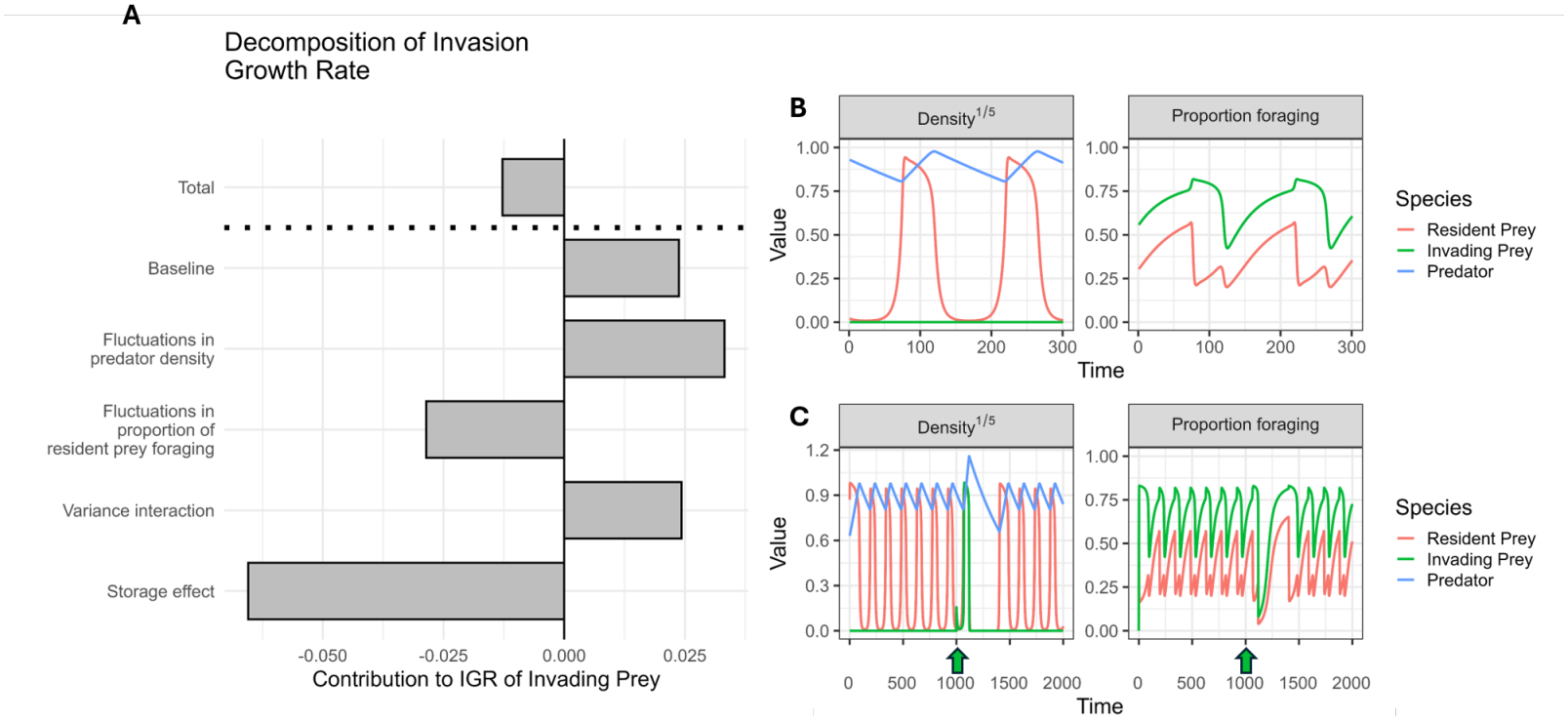
(A) Decomposition of the invasion growth rate of the invading prey species. Individuals bars represent different factors being considered and their respective values represent their contribution to the invasion growth rate. All bars below the dotted line add up to the ‘total’ bar above the dotted line. (B) Time series of the coexistence state in the absence of the invading species. Note that, even in the absence of the invading species, its respective proportion of foragers variable still changes over time. Densities are rescaled to better demonstrate that cyclic dynamics are bounded above extinction. (C) Time series of the attempted invasion of the invading prey species. Upon initial introduction, denoted by the green arrow, dynamics briefly change, following by extinction of the invading species, and consequently converging back to the previous coexistence state. Densities are rescaled to better demonstrate that cyclic dynamics are bounded above extinction.

#### Box 1. Decomposition of invasion growth rate for fluctuation-dependent factors

Here, I briefly describe the decomposition used in Figure 5 with mathematical details provided in Appendix F. We are interested in how fluctuations in predator density (*P*), the proportion of the resident prey population that is foraging (*y*_1_), and their interaction influence the invasion growth rate of the invading prey species relative to the resident prey species. Specifically, I decompose the difference between the invasion growth rates of the invader and resident,

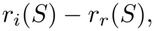

into five components:

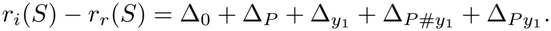

1. **Baseline (**Δ_0_): This term measures the difference in invasion growth rates between the invader and resident when predator density and the proportion of foragers in the resident prey population are fixed at their mean values.
2. **Fluctuations in predator density (**Δ_*P*_ ): This term quantifies the contribution of fluctuations in predator density to the invasion growth rate of the invader relative to the resident. It is calculated by allowing predator density to fluctuate while holding the proportion of foragers in the resident prey population fixed at its mean value, and then comparing the resulting growth rates to the baseline conditions, Δ_0_.
3. **Fluctuations in proportion of resident prey foraging (**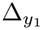): This term quantifies the contribution of fluctuations in the proportion of foragers in the resident prey population to the invasion growth rate of the invader relative to the resident. It is calculated by allowing the proportion of foragers to fluctuate while holding predator density fixed at its mean value, and then comparing the resulting growth rates to the baseline conditions, Δ_0_. Since *y*_1_ appears in the denominator of the predator functional response, fluctuations in *y*_1_ can affect invasion growth rates through Jensen’s inequality. Since the invader and resident can differ in their responses to these fluctuations, 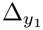 includes contributions of relative nonlinearity.
4. **Variance interaction (**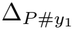): This term quantifies the contribution of interactions between fluctuations in predator density and the proportion of foragers when the two variables are un-correlated. It is calculated by comparing the growth rates obtained from the full fluctuations to those obtained after randomly shuffling the time series of the proportion of foragers. This preserves the marginal distributions of both fluctuating variables while removing their covariance.
5. **Storage effect (**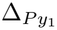): This term quantifies the contribution of the covariance between predator density and the proportion of foragers in the resident prey population. It is calculated as the remaining component of the decomposition after accounting for the baseline, the two fluctuation terms, and the variance interaction term.

## Discussion

By developing a general framework for applying techniques from modern coexistence theory (MCT) to communities of structured populations, I provide a standardized way of identifying when and why invasion or coexistence does (or does not) occur among structured populations. The generality of this framework allows it to be applied across a wide range of ecological contexts, in contrast to previous studies that have often developed similar techniques for applications within a specific subdiscipline (Sieben et al., 2022; Yamamichi et al., 2022; Park et al., 2024). Indeed, the applications explored here span disease ecology, population ecology, and behavioral ecology. Despite the diversity of these systems, this framework successfully calculates invasion growth rates for each model and uses these rates to conduct MCT analyses. More broadly, this framework provides a powerful approach for understanding how population structure influences coexistence and for evaluating how other key ecological factors, such as environmental fluctuations, influence coexistence in communities of structured populations.

While this study emphasizes the broad applicability of the framework, this does not diminish the importance of studies that interpret MCT metrics for specific ecological applications. For instance, Yamamichi et al. (2022) extend niche and fitness differences to models incorporating eco-evolutionary feedbacks. Although the framework presented here can recover the mathematical expressions derived in Yamamichi et al. (2022), the full utility of MCT in an eco-evolutionary context emerges from the biological interpretation of these expressions. By further decomposing niche and fitness differences, Yamamichi et al. (2022) demonstrate that eco-evolutionary niche and fitness differences effectively quantify how niche and fitness differences change between a scenario without evolution and one in which a species has fully evolved in response to its resident competitor. This decomposition allows researchers to identify how ecological and evolutionary processes contribute to coexistence outcomes, highlighting the importance of interpreting MCT metrics in the context of the biological questions being investigated. Similarly, invasion growth rate decompositions can differ depending on which ecological factors are included (Ellner et al., 2019), making it important to select factors that reflect the biological mechanisms of interest. The framework developed here provides the tools needed to derive invasion growth rates and conduct the three core MCT analyses: mutual invasibility, niche and fitness differences, and invasion growth rate decompositions (including the storage effect and relative nonlinearity). However, the broader value of these tools comes from their application to specific ecological systems, where tailoring and interpreting these analyses in the context of biological mechanisms and research questions can generate novel insights into the processes governing coexistence. To this end, this framework is intended not only to facilitate the application of MCT across diverse ecological contexts but also to support discipline-specific studies through which novel ecological insights emerge.

Applying MCT to models with structured population dynamics can reveal ecological insights that may be difficult to identify otherwise. For instance, identifying priority effects is a common motivation for MCT analyses (Ke and Letten, 2018; Grainger et al., 2019; Uricchio et al., 2019), often focused on how initial densities of competitors drive competition exclusion outcomes. However, recent studies suggest that priority effects can depend not only on the initial densities of competing species but also on their initial stage structure (Rasmussen et al., 2014; Zou et al., 2023). Since the methodology proposed here seamlessly incorporates the dynamics of population structure into MCT analyses, we can identify when the impacts of both initial density and population structure on driving coexistence outcomes like priority effects. Indeed, Zou et al. (2023) demonstrated the role of stage structure in driving priority effects and calculated invasion growth rates using a numerical method that leverages the change of coordinates presented here. We demonstrate a similar finding in Figure 3D, where two simulations with identical initial total population densities result in different species being competitively excluded due to differences in initial stage structure. In both simulations, the persisting species begins with five immature and five mature individuals, whereas the competitively excluded species begins with one mature and nine immature individuals. Thus, even when initial total population densities are identical, differences in stage structure (and population structure more generally) can influence competitive outcomes and generate priority effects. This finding highlights the dependence of coexistence outcomes on both total density and population structure, cementing the importance of accounting for within-population variation when investigating the mechanisms underlying coexistence.

Although MCT analyses have only recently been applied to models with explicitly modeled population structure, temporal variation in population structure has long been recognized as a potentially important driver of coexistence, most notably through the storage effect. Perhaps the most well-known example of the storage effect comes from annual plant models in which, within a given year, seeds either remain dormant in a seed bank with some probability of survival or germinate and produce new seeds. Since plants in this system are annual, only seeds are explicitly tracked between years, as all germinated individuals die before the following time step. Nevertheless, species-specific responses of germination rates to environmental fluctuations, together with the covariance between germination rates and the competition experienced by germinated individuals, can generate coexistence that would otherwise be impossible through the storage effect (Chesson, 1994). This example has become so iconic that the storage effect is sometimes misinterpreted as requiring a buffering stage that protects individuals from competition, such as a seed bank, despite this not being a mathematical requirement of the storage effect (Chesson, 1994; Johnson and Hastings, 2022). Here, I demonstrate a storage effect that arises in part from dynamically modeled fluctuations in population structure (Figure 5A). Specifically, predator abundance and the proportion of the resident population foraging negatively covary, generating a negative storage effect for the invading species. Unlike the annual plant model studied by Chesson (1994), where temporal variation arises from environmentally driven variation in germination rates, the fluctuating factor in our model emerges dynamically from changes in population structure. This demonstrates that temporal variation in population structure, whether arising dynamically from population dynamics or through stochastic variation in demographic parameters, can influence invasion and coexistence outcomes through mechanisms such as the storage effect.

Modeling population persistence has long been a central goal of conservation science, with traditional approaches such as population viability analyses (PVAs) emphasizing the role of age and stage structure in determining persistence outcomes (Boyce, 1992; Beissinger, 2002; Ellner et al., 2022a). However, species interactions, which can substantially alter projections of population persistence (Sabo, 2008; Saunders et al., 2018), are often not considered explicitly in these approaches. To address this gap, techniques from modern coexistence theory (MCT) are increasingly applied to conservation and restoration studies to account for species interactions, particularly through invasion growth rate decompositions of fluctuation-dependent mechanisms such as relative nonlinearity and the storage effect (Aoyama et al., 2022; Hallett et al., 2023). Although MCT provides the tools for quantifying how species interactions influence persistence, its applications for restoration and conservation so far have not yet incorporated the role of structured population dynamics. The framework presented here provides a methodological bridge between these approaches, allowing researchers to quantify population growth and persistence while accounting for both population structure and species interactions. Specifically, by combining MCT techniques, such as invasion growth rate decompositions, with traditional PVA analyses, such as sensitivity and elasticity analyses (Easterling et al., 2000; Caswell, 2001; Harper et al., 2011), researchers can disentangle the contributions of population structure and species interactions while also quantifying their possible interactive effects (e.g., the role of increased mortality from gape-limited predators on smaller individuals on population persistence). Combining such analyses with this framework can therefore provide a more holistic assessment of the key ecological drivers of persistence outcomes in at-risk species.

Although I focus here on models with a finite number of classes, the procedure can be extended to populations with a continuous range of classes, such as size-structured integral projection models. Previous studies have calculated the stable size distribution of invading populations by simulating population dynamics over long time periods while constraining total population size to remain small but allowing population structure to change (Chu and Adler, 2015; Ellner et al., 2019). Our approach follows a similar principle but explicitly separates population structure from total population size, allowing the stable size distribution of an invading population to be calculated directly and, consequently, its invasion growth rate. Mathematical details of this extension are provided in Appendix H.

## Conclusion

By providing a general framework for integrating structured population dynamics into modern coexistence theory (MCT), this study enables future research to systematically apply established MCT techniques to understand the mechanisms underlying coexistence in systems with within-species variation. More broadly, this work builds on recent efforts to expand the scope of MCT, contributing to a more generalizable and accessible theoretical framework that can be applied across ecological disciplines.

## Acknowledgements

I thank participants of the Hastings-Schreiber joint lab meeting for thoughtful feedback on an early draft. Additionally, I thank Xuan Kelsy Fei, Brian Lerch, and Chuliang Song for thoughtful written feedback on an early draft. JB acknowledges support from the National Science Foundation Graduate Research Fellowship program under Grant No. DGE-2439024.

## A Bounded dynamics and stability for equilibrium stage structure in exponentially growing population example

Recall the model where the dynamics of a population *x* with stage structure *y* are described by

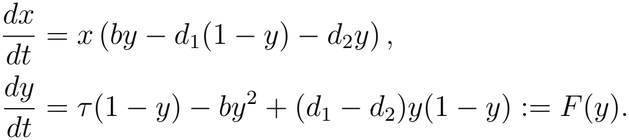

Assume all parameters are strictly positive. Note that *F* (0) = *τ >* 0 and *F* (1) = *−b <* 0. Thus, by the intermediate value theorem, there exists at least one equilibrium *y*^*\**^ *∈* (0, 1) such that *F* (*y*^*\**^) = 0.

Because *F* (*y*) is quadratic, it has at most two roots. Moreover, if it had two distinct roots in (0, 1), then *F* (0) and *F* (1) would have the same sign, since the function would have to cross the horizontal axis twice between 0 and 1. Since *F* (1) *<* 0 *< F* (0), this is impossible. Therefore, the equilibrium *y*^*\**^ *∈* (0, 1) is unique.

Finally, because *F* (*y*) *>* 0 immediately left of *y*^*\**^ and *F* (*y*) *<* 0 immediately right of *y*^*\**^, trajectories flow toward *y*^*\**^ from both sides. Therefore, *y*^*\**^ is locally stable.

## B Leading eigenvalue is equivalent to invasion growth rate in exponentially growing population with stage structure example

Recall

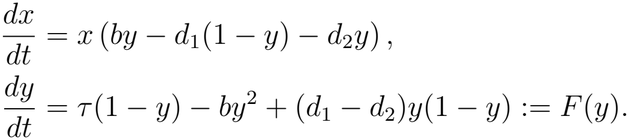

from Appendix A.

### B.1 Solving for biologically relevant *y*^*\**^

I demonstrate in Appendix A that the equilibrium *y*^*\**^ *∈* (0, 1) exists and is unique. Solving explicitly for *y*^*\**^, I find that

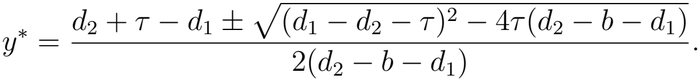

To determine the biologically relevant root, we assume all parameters are strictly positive and assess two cases: when *d*_2_ *− b − d*_1_ *>* 0 and when *d*_2_ *− b − d*_1_ *<* 0.

Let *A* = *d*_2_*−b−d*_1_ and *B* = *d*_1_*−d*_2_*−τ*, so that the equilibrium equation is *Ay*^2^+*By*+*τ* = 0, with roots

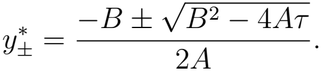

If *A <* 0, then

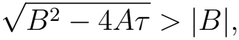

so 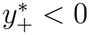 and 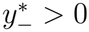 . Thus 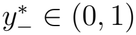

If *A >* 0, then *d*_2_ *> b* + *d*_1_. Therefore, *B* = *d*_1_ *− d*_2_ *− τ < d*_1_ *−* (*b* + *d*_1_) *− τ* = *−b − τ <* 0. Thus, *B <* 0, meaning both roots are positive. Since *F* (1) *<* 0 *< F* (0) and the positive root is unique in (0, 1), we have 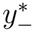 *∈* (0, 1). Thus, *y*^*\**^ is again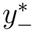.

Therefore, we consider the biologically relevant root

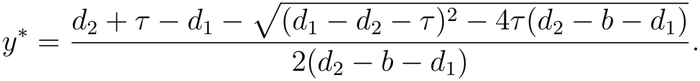

when *A≠*0. If *A* = 0, then 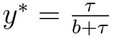.

### B.2 Invasion growth rate of transformed equations

Recall

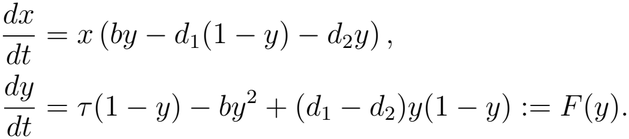

Since we wish to calculate the invasion growth rate of species *x*, we evaluate its per-capita growth rate when *x* = 0 and *y* = *y*^*\**^. This gives

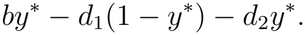

### B.3 Leading eigenvalue of matrix model

Recall

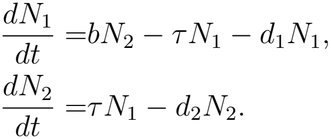

where *N* = *x*(1 *− y*) and *N*_2_ = *xy*. If we let *N* = (*N* _1_, *N* _2_)^*T*^ and 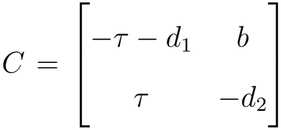, then we have

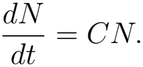

Solving for the eigenvalues of *C*, we find

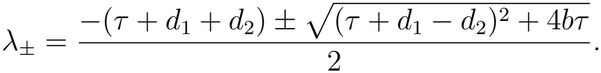

Thus, the leading eigenvalue is

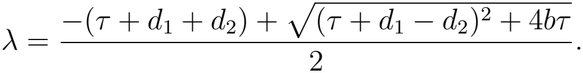

### B.4 Demonstrate equivalence of leading eigenvalue of matrix model and invasion growth rate of transformed model

Recall the leading eigenvalue of the matrix model is

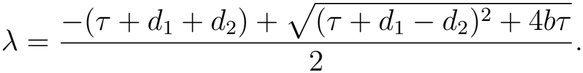

and the invasion growth rate of species *x* = *N*_1_ + *N*_2_ is

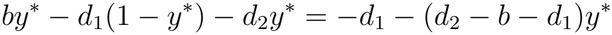

Where

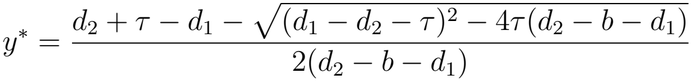

when *d*_2_ *− b − d*_1_*≠*0.

We can simplify *y*^*\**^ such that

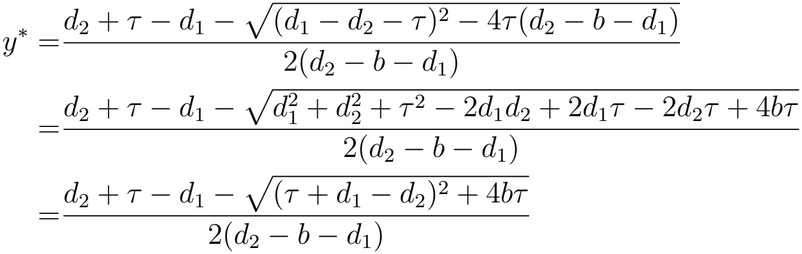

Putting this value into the invasion growth rate, we have

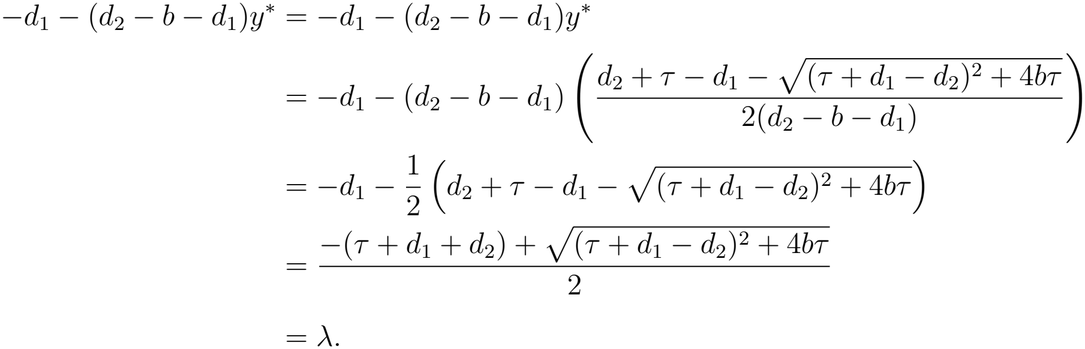

Thus, the leading eigenvalue of the matrix model is equivalent to the invasion growth rate of the transformed model, as proven more generally in Patel and Schreiber (2018).

## C Parameters for disease model

**Table 1:** Parameterization for model of two strains of influenza with cross-immunity.

| ***H1N1*** |  |  |
|---|---|---|
| Parameter | Description | Value |
| μ | Birth rate into susceptible population | 1/(80 × 52) |
| δ<sub>1</sub> | Waning immunity rate | 1/(3.12 × 52) |
| σ<sub>11</sub> | Cross-immunity from H1N1 to H1N1 | 1 |
| σ<sub>12</sub> | Cross immunity from H3N2 to H1N1 | 0.17 (varies) |
| β<sub>1</sub> | Transmission rate | 1.44 × (μ + γ<sub>1</sub>) |
| γ<sub>1</sub> | Recovery rate | 7/2.64 |
| ***H3N2*** |  |  |
| Parameter | Description | Value |
| μ | Birth rate into susceptible population | 1/(80 × 52) |
| δ<sub>2</sub> | Waning immunity rate | 1/(3.08 × 52) |
| σ<sub>22</sub> | Cross-immunity from H3N2 to H3N2 | 1 |
| σ<sub>21</sub> | Cross immunity from H1N1 to H3N2 | 0.40 |
| β<sub>2</sub> | Transmission rate | 1.60 × (μ + γ<sub>2</sub>) |
| γ<sub>2</sub> | Recovery rate | 7/3.03 |

## D Parameters for perennial plant model

**Table 2:**
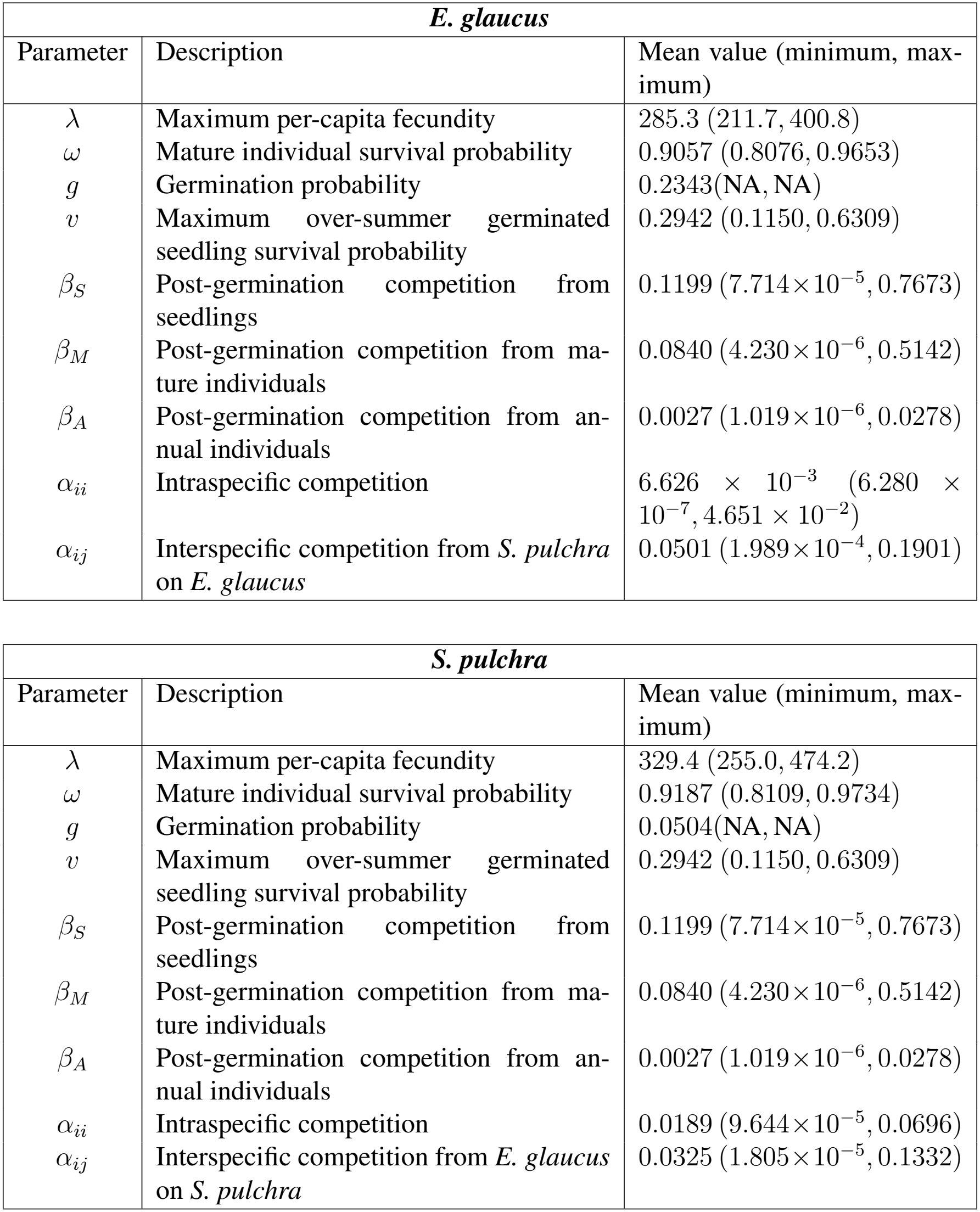
Parameterization for model of competition between perennial plants.

| ***E. glaucus*** |  |  |
|---|---|---|
| Parameter | Description | Mean value (minimum, max-imum) |
| λ | Maximum per-capita fecundity | 285.3 (211.7, 400.8) |
| ω | Mature individual survival probability | 0.9057 (0.8076, 0.9653) |
| *g* | Germination probability | 0.2343(NA, NA) |
| *v* | Maximum over-summer germinated seedling survival probability | 0.2942 (0.1150, 0.6309) |
| β<sub>*S*</sub> | Post-germination competition from seedlings | 0.1199 (7.714×10<sup>−5</sup>, 0.7673) |
| β<sub>*M*</sub> | Post-germination competition from ma-ture individuals | 0.0840 (4.230×10<sup>−6</sup>, 0.5142) |
| β<sub>*A*</sub> | Post-germination competition from an-nual individuals | 0.0027 (1.019×10<sup>−6</sup>, 0.0278) |
| α<sub>*ii*</sub> | Intraspecific competition | 6.626 × 10<sup>−3</sup> (6.280 × 10<sup>−7</sup>, 4.651 × 10<sup>−2</sup>) |
| α<sub>*ij*</sub> | Interspecific competition from *S. pulchra* on *E. glaucus* | 0.0501 (1.989×10<sup>−4</sup>, 0.1901) |
| ***S. pulchra*** |  |  |
| Parameter | Description | Mean value (minimum, max-imum) |
| λ | Maximum per-capita fecundity | 329.4 (255.0, 474.2) |
| ω | Mature individual survival probability | 0.9187 (0.8109, 0.9734) |
| *g* | Germination probability | 0.0504(NA, NA) |
| *v* | Maximum over-summer germinated seedling survival probability | 0.2942 (0.1150, 0.6309) |
| β<sub>*S*</sub> | Post-germination competition from seedlings | 0.1199 (7.714×10<sup>−5</sup>, 0.7673) |
| β<sub>*M*</sub> | Post-germination competition from ma-ture individuals | 0.0840 (4.230×10<sup>−6</sup>, 0.5142) |
| β<sub>*A*</sub> | Post-germination competition from an-nual individuals | 0.0027 (1.019×10<sup>−6</sup>, 0.0278) |
| α<sub>*ii*</sub> | Intraspecific competition | 0.0189 (9.644×10<sup>−5</sup>, 0.0696) |
| α<sub>*ij*</sub> | Interspecific competition from *E. glaucus* on *S. pulchra* | 0.0325 (1.805×10<sup>−5</sup>, 0.1332) |

## E Removing perenniality from perennial plant model

### One missing perenniality

Let species 1 be the species with perenniality and species 2 be the species without. The model is then

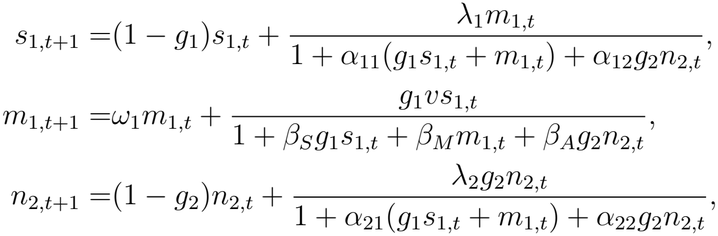

where *β*_*A*_ is the impact of competition from a species without stage structure. Let *n*_1,*t*_ = *s*_1,*t*_ + *m*_1,*t*_ and 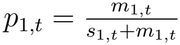. Then, I have

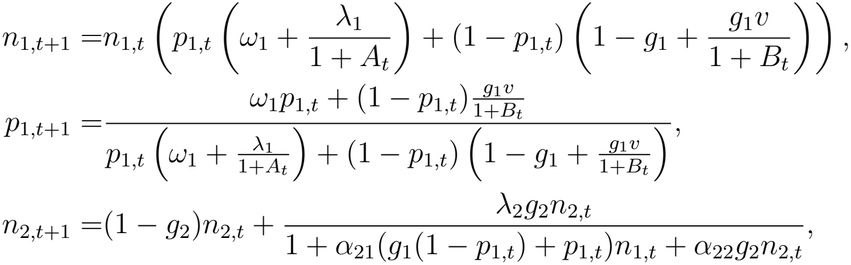

Where 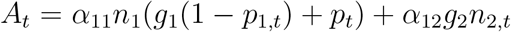 and 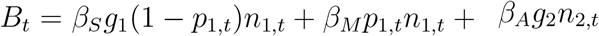

### Neither with perenniality

If neither species has perenniality, I use the following model for both species:

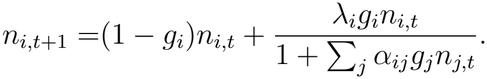

## F Decomposition of invasion growth rate

Here, I explain the decomposition performed in the manuscript based on the technique proposed by Ellner et al. (2019) using notational language from Brennan and Schreiber (2026). Say we are interested how some set of ecological factors impact the invasion growth rate of species *i* at coexistence state *S*. Specifically, we are interested in how fluctuations in predator abundance (*θ*_1_) and proportion of foragers in the resident prey population (*θ*_2_) impact invasion success. Let the invasion growth rate of species *i* for the ecological factors of interest *θ* = (*θ*_1_, *θ*_2_) be *r*_*i*_(*S, θ*) where *θ*_*ℓ*_ = 0 denotes factor *ℓ* being fixed at its average while *θ*_*ℓ*_ = 1 denotes fluctuations in the respective factor. Then, the invasion growth rate of species *i* at coexistence state *S* can be written as:

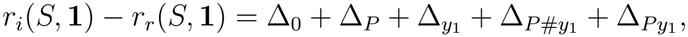

where **1** = (1, 1). Δ_0_ denotes the baseline value of the invader’s growth rate relative to the resident’s growth rate when neither factor is fluctuating. Specifically,

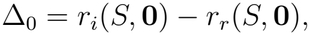

where **0** = (0, 0) and *r* is the resident prey population.

Δ_*P*_ and 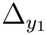 denotes the impact of fluctuations of the respective factor on the invasion growth rate of the invading prey population relative to the resident prey population. Specifically,

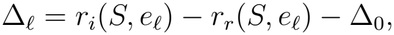

where *e*_*ℓ*_ is the vector with 1 in entry *ℓ* and 0 elsewhere.

The total interaction term, denoted Δ_*L*_ is then given by

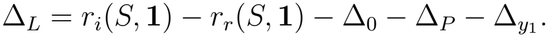

I then decompose this term into two terms, 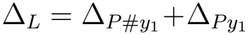 . Let *S*^#^ denote the coexistence state *S* but with the timeseries tracking *y*_1_ shuffled, preserving its marginal distribution but ridding of correlation between *y*_1_ and *P* . Then, I have

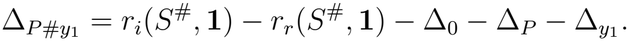

Lastly, I have

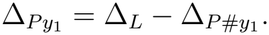

## G Parameterization for model of foraging-predation risk trade-off

**Table 3:** Parameterization for model of foraging-predation risk tradeoff.

| ***Resident Prey*** |  |  |
|---|---|---|
| Parameter | Description | Value |
| *r* | Intrinsic growth rate when foraging | 3 |
| *a*<sub>*IP*</sub> | Predator attack rate when resident is in idle class | 1 |
| *a*<sub>*FP*</sub> | Predator attack rate when resident is in foraging class | 4 |
| *τ*<sub>*FI*</sub> | Average time spent in idle class | 6 |
| *τ*<sub>*IF*</sub> | Average time spent in foraging class | 1 |
| ***Invading Prey*** |  |  |
| Parameter | Description | Value |
| *r* | Intrinsic growth rate when foraging | 2.64 |
| *a*<sub>*IP*</sub> | Predator attack rate when invader is in idle class | 2 |
| *a*<sub>*FP*</sub> | Predator attack rate when invader is in foraging class | 5 |
| *τ*<sub>*FI*</sub> | Average time spent in idle class | 3 |
| *τ*<sub>*IF*</sub> | Average time spent in foraging class | 2 |
| ***Predator*** |  |  |
| Parameter | Description | Value |
| *c* | Conversion rate | 0.1 |
| *μ* | Predator death rate | 0.01 |
| *h* | Handling time of predator | 2 |

## H Change of variables for an integral projection model

Let *n*_*i*_(*z*_*i*_, *t*) denote the density of individuals of species *i* with size *z*_*i*_ at time *t*. Thus, the total number of individuals of species *i* is given by

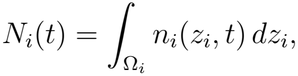

where Ω_*i*_ gives the possible range of *z*_*i*_. Let 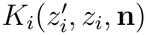 denote the integral projection kernel for species *i*, describing the contribution of individuals of size *z*_*i*_ at time *t* to the expected number of individuals of size 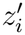 at time *t* + 1 in the presence of the community

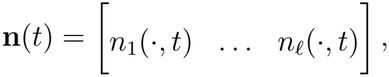

consisting of *ℓ* potentially size-structured species. Note that the kernel may depend on the abundance and size distribution of every species in the community. Thus, individuals of species *i* may experience demographic effects that depend not only on the abundance of other species, but also on their size distributions. The model for species *i* can then be written as

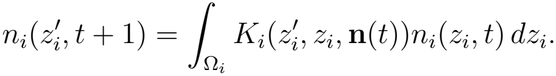

We now perform a change of variables. Let

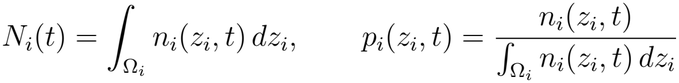

where *N*_*i*_(*t*) denotes the total population size of species *i* and *p*_*i*_(*z*_*i*_, *t*) denotes the size distribution of species *i*. Thus,

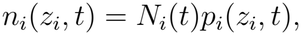

and

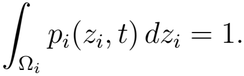

Let

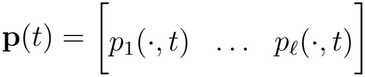

denote the matrix of size distributions across all *ℓ* species. Let *N* (*t*) denote the vector of total population sizes across the community. Because

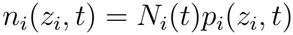

for each species, we can equivalently express the kernel as a function of **p**(*t*) and *N* (*t*). Specifically, let

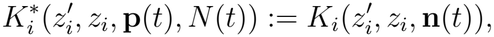

where 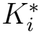 is the kernel expressed as a function of the population sizes and size distributions rather than the population densities directly.

The total population at time *t* + 1 is then given by

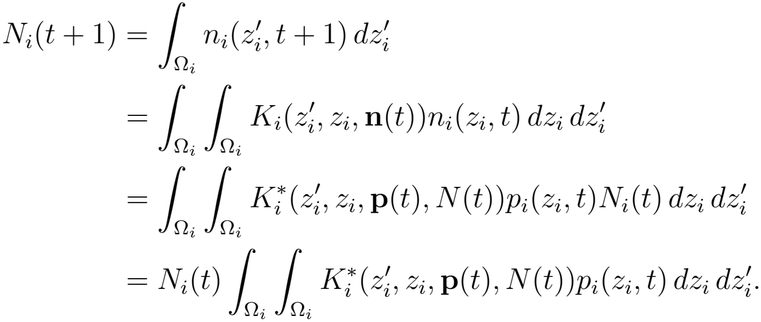

Similarly, the transformed size distribution satisfies

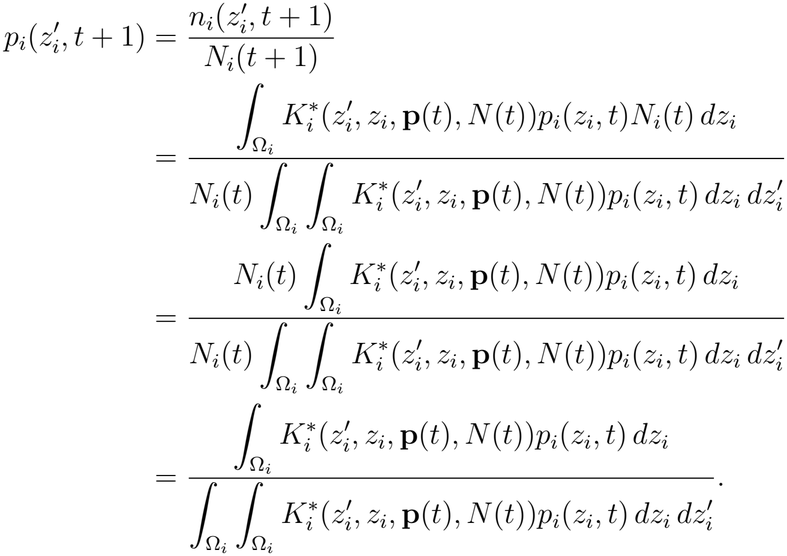

Thus, the transformed system can be written as

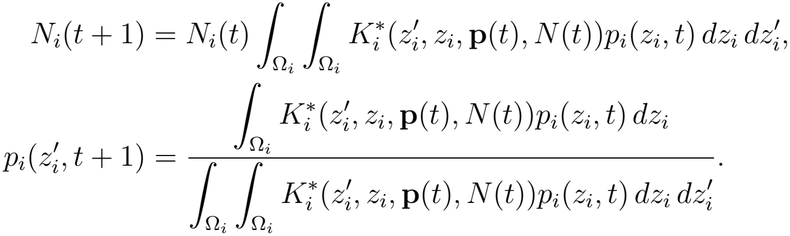

We can now consider the invasion growth rate of species *i*. Specifically, we allow the resident community, with *N*_*i*_(*t*) = 0, to reach its asymptotic dynamics. Let 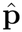 denote the stable size distributions of the resident species and 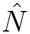 their corresponding equilibrium abundances.

Then, invasion growth rate at coexistence state *S* is

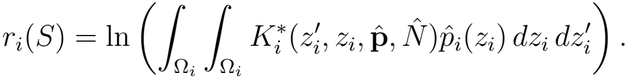

